# Genomic Prediction Enables Provenance-Aware Selection in Sessile Oak (*Quercus petraea*) using Foliar Physiological Traits

**DOI:** 10.64898/2026.03.31.715316

**Authors:** Leke Victor Aiyesa, Miao He, Henning Wildhagen, Wilfried Steiner, Andre Hardtke, Martin Hofmann, Markus Mueller, Oliver Gailing

## Abstract

Climate change is reshaping the adaptive landscape of forest ecosystems, demanding more efficient strategies to identify and deploy resilient tree genotypes. Genomic prediction offers a powerful framework to accelerate selection for complex physiological traits underlying climate adaptability in long-lived species such as sessile oak (*Quercus petraea* (Matt.) Liebl.).

Here, we conducted genomic prediction for three key physiological traits carbon isotope composition, nitrogen isotope composition, and the carbon-to-nitrogen content ratio (C/N ratio) measured in 746 trees genotyped with dense genome-wide markers (∼580,000 SNPs). High genomic heritabilities were estimated across traits, with within-year prediction accuracies (Pearson’s *r* between genomic estimated values and observed phenotypes) reaching 0.77. Notably, across-year and across-provenance predictions remained substantial (0.41 < r < 0.82), with predictability declining with increasing genetic distance (*F_ST_*) between training and test provenances for nitrogen isotope composition and C/N ratio. In addition, GWAS-guided SNP preselection increased heritability capture by ∼15% relative to random SNP subsets. And, the pronounced provenance-by-environment interactions observed indicated substantial phenotypic plasticity in these traits.

These results demonstrate the strong potential of applying genomic prediction to foliar physiological traits as polygenic predictors for climate adaptation in plants, support provenance-aware breeding to improve forest establishment, and provide practical strategies for deploying genomic prediction in long-lived species.

## Introduction

The application of prediction-based breeding in plants has advanced considerably over the past two decades, particularly for the early identification of candidates for selection and for complex polygenic traits (Crossa et al., 2017; Alemu et al., 2024). However, prediction accuracy tends to decline as the genetic and/or geographic divergence between the training and target populations increases, thereby reducing the consistency of model performance across populations and diverse geographical origins (Crossa et al., 2017). Empirical studies examining how genetic differentiation impacts prediction accuracy (Rolling et al., 2024; Vitale et al., 2025) have been conducted primarily for crop species. Very few, if any, studies have considered this effect in forest tree species within a genomic prediction framework, despite growing recognition of its importance in forest tree breeding programs (Li et al., 2017; Aspalter et al., 2025). Meanwhile, the mechanisms underlying these clinal genetic interactions particularly for polygenic traits remain poorly understood (Alemu et al., 2026; Kremer et al., 2025; Lenth & Piaskowski, 2017), including adaptive phenotypic plasticity (Brendel & Epron, 2021; Kurjak et al., 2024).

Exploiting genetic components within a genomic prediction framework for polygenic traits requires dense genotyping because linkage disequilibrium decays rapidly across the genome, particularly in forest trees (Ingvarsson, 2005; de Souza et al., 2018; Grattapaglia et al., 2018). Although high-throughput sequencing now produces massive SNP datasets, analytical bottlenecks remain in processing this data deluge. Moreover, including all SNP markers indiscriminately does not necessarily improve prediction accuracy and can substantially increase computational demands (Resende et al., 2012; Isik, 2016). Consequently, analytical approaches such as GWAS-based SNP preselection have been used to manage large marker sets for downstream genomic prediction analyses (Chen et al., 2023; Zhang et al., 2023; El-Kassaby et al., 2024; Munyengwa et al., 2025). In this approach, informative markers are typically preselected based on their association strength and then used in subsequent prediction analyses.

Existing genomic prediction studies in forest trees have primarily focused on species such as spruce, pine, and eucalypts, targeting traits related to growth, wood quality, disease resistance, and morphology—including height, volume, and diameter (Thistlethwaite et al., 2020; Akutsu et al., 2023; Duarte et al., 2024; El-Kassaby et al., 2024; Slavov et al., 2025). These studies have demonstrated the feasibility and promise of genomic prediction for accelerating genetic gain in long-lived tree species (Isik, 2014; Grattapaglia, 2022). So far, only a few genomic prediction studies have been conducted in major forest tree species such as oak (Degen et al., 2025), and none have yet leveraged whole-genome sequencing datasets to model foliar physiological traits indicative of climatic adaptation for this species.

*Quercus petraea* (Matt.) Liebl. (sessile oak) is native to much of Europe and parts of the Middle East and is characterized by high genetic diversity, potentially enabling the species to adapt to rapidly changing environmental conditions (Thomas et al., 2002; Härdtle et al., 2014). Over the past decade, the *Quercus* genus has experienced a remarkable expansion in genomic research, driven by advances in high-throughput sequencing technologies (Sork et al., 2022). Landmark efforts have focused on assembling high-quality reference genomes for key species (*Quercus robur* L., Plomion et al., 2018; *Quercus alba* L., Larson et al., 2024), providing a genomic framework for identifying genome-wide polymorphisms essential for understanding genetic diversity (Leroy et al., 2019; Nocchi et al., 2021) and evolutionary patterns (Plomion et al., 2018). Building on these resources, population-scale sequence data have helped in dissecting the genetic architecture of complex traits via GWAS in both natural and breeding populations (Saleh et al., 2022; Leroy et al., 2019). Many of these traits, including physiological traits, are polygenic, making them difficult to dissect using GWAS alone and to predict (Holliday et al., 2015; Cortés et al., 2020; Sharma et al., 2023).

Foliar physiological traits such as carbon isotope composition (δ¹³C), nitrogen isotope composition (δ¹⁵N), and the leaf carbon-to-nitrogen ratio (C/N ratio) have been widely reported as proxies for intrinsic water-use efficiency (iWUE), nutrient acquisition efficiency, and drought- and nutrient-stress responses in forest trees (Nguyentu et al., 2004; Yang et al., 2015; Doby et al., 2024; Nosenko et al., 2025). These physiological traits also exhibit strong relationships with growth-related traits such as leaf area, leaf thickness, and leaf nitrogen content (Cumbie et al., 2011; Famula et al., 2018). δ¹³C reflects the ratio of ¹³C to ¹²C in leaf tissues and serves as an integrative measure of iWUE, capturing the trade-off between stomatal conductance and photosynthetic carbon assimilation (Bartholomé et al., 2015; Zhao et al., 2019). Variation in δ¹³C is frequently associated with drought tolerance and habitat water availability (Brendel et al., 2007), and has been widely used to assess within-species genetic variation in iWUE in long-lived tree species (e.g. Brendel et al., 2007; Bogeat-Triboulot et al., 2019; Nosenko et al., 2025). Studies have further reported significant intraspecific variation in δ¹³C and δ¹⁵N among oak populations along gradients of water and nitrogen availability (Spasojevic & Weber, 2021; Nosenko et al., 2025). Moreover, moderate-to-high heritability estimates for these traits, ranging from 0.21 to 0.72 (McKown et al., 2013; Alexandre et al., 2020), together with association analyses (Famula et al., 2018; Tost et al., 2025), support their polygenic control and their suitability as targets for genomic selection in breeding programmes for climate-resilient forests.

In this study, we implemented genomic prediction for δ¹³C, δ¹⁵N, and the C/N ratio using ∼580,000 SNP markers derived from targeted reduced-representation sequencing of 746 sessile oak trees, representing eight provenances established at two sites in Germany. Using a GWAS-guided SNP selection approach, we evaluated the performance of prediction models in estimating trait genomic heritability and, ultimately, their predictive ability under practicable tree breeding scenarios. We hypothesized that these traits are suitable targets for genomic selection for climate adaptation in oaks, and that the high-density marker set would capture a large proportion of the genomic regions underlying these traits, thereby improving prediction performance, particularly in out-of-provenance prediction scenarios.

## Materials and Methods

### Plant Material and Experimental Design

A total of 746 trees from eight provenances across four European countries (France: 185; Denmark: 92; United Kingdom: 280; Germany: 189) were sampled for phenotyping and genome-wide genotyping. The study was conducted at two provenance trial sites in Germany—Harz (latitude: 51°36′37″ N, longitude: 10°38′32″ E, elevation: 372 m.a.s.l.) and Unterlüß (latitude: 52°46′34″ N, longitude: 10°28′14″ E, elevation: 117 m a.s.l.) managed by the Northwest German Forest Research Institute (NW-FVA). The provenances and sites used here are part of the international “Madsen Collection” (Madsen, 1990), a coordinated provenance trial series established in 1992–1993 across 27 sites with seedlots from 19 *Quercus petraea* provenances covering the species’ entire natural range (see Madsen, 1990 and Sáenz-Romero et al., 2017 for further details).

At the site Harz, the trial was arranged as a Randomized Incomplete Block Design, while Randomized Complete Block Design was used at site Unterlüß, both with three replicates. Each replicate includes one plot comprising all 19 provenances used in the experiment. Sampling took place over two consecutive years: between July and August 2021 and again in June 2022. A standardized sampling protocol was applied to ensure consistency across sites and replicates. Detailed environmental conditions recorded during sampling are presented in Supplementary Table 4, while geographic and environmental characteristics of the provenances are shown in Figure 1 and Supplementary Table 1.

**Figure 1.**
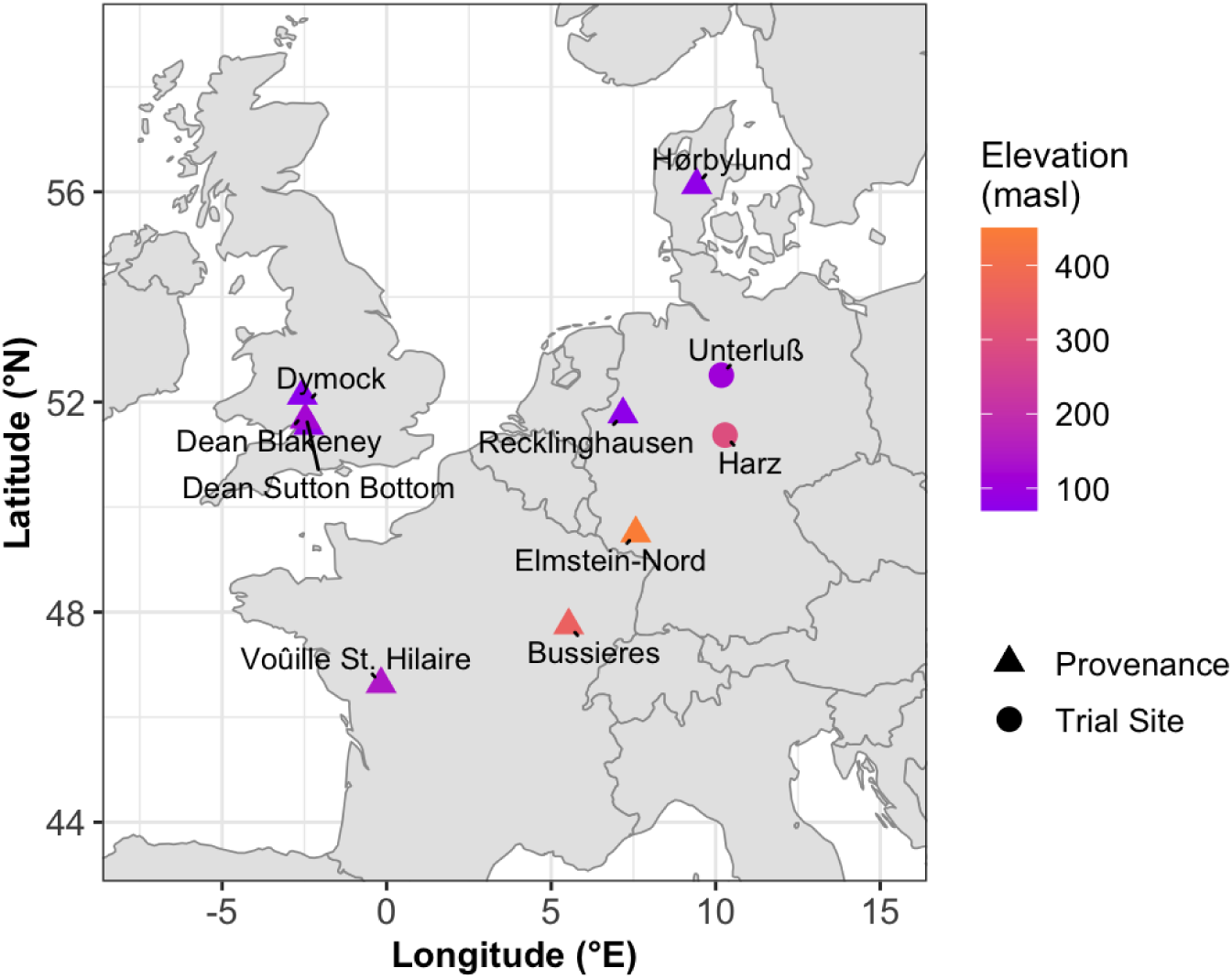
Geographical distribution of provenances used in this study showing their latitude, longitude and elevation. Circle indicate the trial sites in Germany where trees were sampled.

### Sample Preparation and Isotope Analysis

Fresh leaf samples were collected and oven-dried at 60 °C until they were completely desiccated. The dried plant material was then transferred to 2 ml microcentrifuge tubes containing three stainless steel beads and homogenized using a mixer mill (Retsch MM 300 TissueLyser) for 2 minutes. From the homogenized powder, subsamples of 1.0-1.5 mg were precisely weighed into tin capsules. These capsules were arranged in 96-well trays and sealed by compression in preparation for stable isotope analysis. We investigated three key physiological traits: carbon isotope composition (δ¹³C), nitrogen isotope composition (δ¹⁵N), and the carbon-to-nitrogen content ratio (C/N ratio). The analysis was performed at the Center for Stable Isotope Research and Analysis at the University of Göttingen. The homogenized plant material was analyzed using an elemental analyzer coupled with an isotope ratio mass spectrometer (IRMS) to determine δ¹³C and δ¹⁵N values. The C/N ratio was calculated from the molar content (mmol/g) and the atomic weights of carbon and nitrogen.

### Phenotypic Trait Analysis

Phenotypic differences among provenances were evaluated for δ¹³C, δ¹⁵N, and the C/N ratio. We applied a factorial design to assess (1) differences among the eight provenances within the same site and year, (2) differences between the two sites within the same year, and (3) differences between years within the same site. Prior to analysis, all trait data were tested for normality and homogeneity of variances using the Shapiro–Wilk and Bartlett tests, respectively. For δ¹³C, which approximated a normal distribution, we fitted linear mixed-effects models using the lme4 R package (Bates et al., 2015). Provenance, site, and year were treated as fixed effects, with individual trees modeled as a random effect. For δ¹⁵N and the C/N ratio, which deviated from normality, we fitted generalized linear mixed-effects models. Post hoc pairwise comparisons among groups were performed using the emmeans R package (version 1.8.6; Lenth, 2017) with a Tukey adjustment for multiple testing. All analyses were conducted in R version 4.4.0, and statistical significance was evaluated at an α level of 0.05.

### Genotyping and Variant-Calling

DNA was isolated from leaf material using the Dneasy Plant Min Kit (Qiagen), and genotyping was performed using Single Primer Enrichment Technology (SPET), a targeted genotyping by sequencing (GBS) approach widely applied in forest trees, crops, and other non-model species (Scaglione, Pinosio et al. 2019). Sequenced reads were aligned to the oak (*Quercus robur*) reference genome version 2 (Plomion et al., 2018) using BWA-MEM v0.7.17 (Li and Durbin, 2009). Reads with a mapping quality score below 10 were discarded, and duplicate reads were subsequently removed to ensure data quality. Variants were filtered to retain only sites with minor allele frequency (MAF) ≥ 0.01, per-variant missingness ≤ 10%, site quality score (QUAL) ≥ 30, and genotype read depth between 10 and 50, using VCFtools v0.1.16 (Danecek et al., 2011). Genotype imputation was performed with Beagle (v5.x) (Browning et al., 2018) using the study sample set. Further filtering removed sites with Indels, monomorphic sites, and variants with alleles discordant with the reference panel, leaving the remaining sites for downstream analysis.

### GWAS and SNP-Selection

GWAS was conducted prior to genomic prediction, using 90% of individuals randomly sampled to mimic a training set in a 10-fold cross-validation scheme. The Bayesian-information and Linkage-disequilibrium Iteratively Nested Keyway (BLINK) model, as implemented in the GAPIT R package version 3.0 (Huang et al., 2018), was used for association testing. The model can be expressed as:

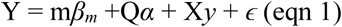

Where Y is the vector of phenotypic values, *β_m_* is the fixed effect of the marker being tested, m is the vector of the genotypes of the marker being tested, Q is a matrix of population structure covariates (accounted for with the first two principal components and site information), and α is the vector of effects for the population structure. X is a matrix of other significant markers fitted as fixed covariates, with *y* as the vector of their effects, *ɛ* is the vector of residual errors, typically assumed to be *ɛ ∼ N*(0,*σ_e_^2^I*).

To select informative SNPs for prediction, markers were chosen at different p-value thresholds (10⁻⁵, 10^−4^, 10⁻³, 10⁻², 0.05, 0.1, 0.5, 0.8). These SNP subsets were subsequently used in a GBLUP framework to estimate the proportion of additive genetic variance explained (genomic heritability (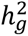)). The threshold beyond which the inclusion of additional SNPs no longer substantially increased additive variance was considered optimal for genomic prediction.

### Genomic Prediction Models

Three genomic prediction models, Genomic Best Linear Unbiased Prediction (GBLUP, VanRaden 2008), the Bayesian Ridge Regression (BRR, Pérez and de los Campos, 2014), and the Light Gradient Boosting (LGM; Shi et al., 2020) were used based on previous reports on their suitability for GP modeling. These three models were fitted for each year and for each trait. The GBLUP model leverages a whole-genome relationship matrix to capture polygenic effects, which are explained by the relationship among the individuals present in the dataset, given as:

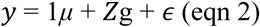

where *y* is an *n*-dimensional vector of phenotypes for the trait, 1 represents an n-dimensional vector of ones, and denotes the overall mean of the set. *Z* represents the genomic random effects, and *ε* is the vector of residual error. *Z* and ɛ are considered as random parameters having multivariate normal distribution *g*∼*N*(0,*G*) and *ε*∼*N*(0,*G*_*e*_), respectively. The covariance matrix *Z* is modeled by the additive genomic relationship matrix, which was computed following the first method described by VanRaden (2008).

The BRR model, a shrinkage-based Bayesian method that assumes equal variance across markers, was implemented following:

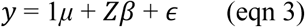

where *Z* is the genotypic matrix and *β* is the vector of marker effects. All other notations are the same as mentioned in the GBLUP model. The marker effects are independently and identically distributed under the Gaussian assumption with the same variance for all the effects, i.e. 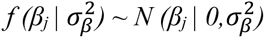, and the overall mean μ is assigned a flat prior. The variance of marker effects and the error variance are assumed to follow an inverse chi-squared distribution (χ⁻²(v, S)), where v represents the degrees of freedom, and S is the scale parameter. The prior distribution for the scale parameter S is modeled using a Gamma distribution with a rate parameter (r) and shape parameter(s). The GBLUP and BRR methods were implemented using the BGLR R package (Pérez and de los Campos, 2014).

The machine learning algorithm LightGBM (LGB), a tree-based gradient boosting framework capable of modelling non-linear relationships and interactions (Alemu et al., 2024), was applied for genomic prediction. For each trait, we trained an LGB regression model using 1,000 boosting iterations (nrounds = 1000), specifying a regression objective using Pearson’s correlation as evaluation metric. The learning rate was set to 0.1, the maximum tree depth to 6, the maximum number of leaves per tree to 31, and subsampling was applied to both features and instances, with the bagging fraction fixed at 1.0. All remaining hyperparameters were left at their default settings as implemented in the R package lightgbm, version 4.3.0 (Shi et al., 2020).

### Model Evaluation and Cross-Validation Strategy

Model performance was assessed using the Pearson correlation coefficient (r) between observed phenotypic values and genomic estimated breeding values (GEBVs) in a 10-fold cross-validation framework. In the within-year genomic prediction scenario, individuals were randomly partitioned into training (90%) and test (10%) sets in each fold. To evaluate model robustness under increasingly challenging conditions, we additionally implemented two provenance-based validation schemes. First, in a leave-one-provenance-out (within-year) scheme, one of the eight provenances was excluded from the training set in each fold and used exclusively for testing, while all remaining provenances served as the training set; this scenario assesses the model’s ability to predict across genetically distinct provenances within the same year and sites. Second, in a leave-one-provenance-and-one-year-out scheme, all observations from a given provenance for a specific year were excluded from the training dataset and used only for testing; here, the model was trained on data from the remaining seven provenances for other year and evaluated on the excluded provenance, representing a more stringent validation that approximates real-world deployment where predictions are required for novel provenance–environment–year combinations.

### Genetic Differentiation and Prediction Ability

To assess the impact of genetic divergence on prediction accuracy, fixation indices (*F_ST_*) were calculated between the test provenance and the training provenances in the leave-one-provenance-and-one-year-out cross-validation scenario. *F_ST_* values were estimated using the hierfstat R package, version 0.5 (Goudet, 2004), which implements Wright’s F-statistics for hierarchical population structure. Pairwise *F_ST_* was computed based on the GWAS pre-selected SNPs for each trait, and thus reflects the degree of genetic differentiation between the excluded (test) provenance and all of the remaining (training) provenances. The relationship between prediction ability, measured as the Pearson correlation (r) between observed phenotypes and genomic estimated values, and *F_ST_*, was evaluated using linear regressions. This analysis aimed to determine whether increasing genetic divergence between training and test populations was associated with a decline in prediction accuracy, thereby informing the limits of genomic prediction transferability across divergent provenances in sessile oak (*Quercus petraea*).

## Results

### Phenotype Data Analysis

A strong, positive correlation between years 2021 and 2022 was observed for δ¹³C and δ¹⁵N (Supplementary Figure 1), indicating high repeatability of these traits, particularly at the Unterlüß site compared to Harz. In contrast, the C/N ratio showed weaker year-to-year correlation, suggesting that it is more sensitive to year-specific environmental conditions, such as differences in temperature, moisture, or nutrient availability.

Clear spatial heterogeneity in trait expression was observed across sites (Harz vs. Unterlüß) and years (Supplementary Figure 2). Within each year, trait distributions frequently differed significantly between sites for δ¹³C, δ¹⁵N, C/N ratio. This difference was observed for all provenances for δ¹³C suggesting site-specific effects on carbon assimilation or water-use patterns. Provenance Horbylund (Denmark) consistently showed significant site-specific effects across the three traits and for the two years. δ¹⁵N displayed greater variability in Harz, potentially reflecting differences in nitrogen availability, soil processes, or uptake efficiency between sites. This provenance-level differentiation, combined with site-specific contrasts, reveals significant provenance-by-environment interactions (PEI) for all three traits that were consistent across years for δ¹⁵N (Supplementary Figure 3 and Table 3). These results underscore the importance of accounting for genetic origin in genomic selection for *Quercus* species, particularly for δ¹⁵N.

### Genomic Heritability

To assess the heritability and polygenic architecture of the traits, genomic heritability (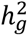) was estimated using SNP sets selected under progressively relaxed GWAS P-value thresholds. For δ¹³C, δ¹⁵N, and the C/N ratio, 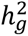 increased progressively as the number of SNPs increased until a threshold of P < 0.05 was reached. After this, the gain only marginally increased and diminished at P<0.5 with about 250,000 SNPs (**Figure 2**). At P < 0.05, SNPs were evenly distributed within and across all twelve *Quercus robur* chromosomes for each trait (Supplementary Figure 4). Using the GWAS-guided SNP selection approach, peak 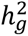 estimates were higher for δ¹⁵N (0.78) and the C/N ratio (0.76) than for δ¹³C (0.72). Meanwhile, at stringent P-value thresholds, 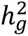 was also low, confirming the polygenicity of these traits. GWAS-guided SNP selection yielded higher 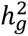 than random selection of SNPs, particularly at more stringent p-value thresholds. Moreover, the overall pattern of increase or decline in 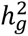 was consistent in the GWAS-guided SNP selection approach and the random SNP selection approach, indicating that GWAS-based SNP preselection is robust and reliable for identifying SNP-markers to be deployed for genomic prediction. The number of SNPs retained at each p-value threshold for each trait are shown in Supplementary Figure 5.

**Figure 2.**
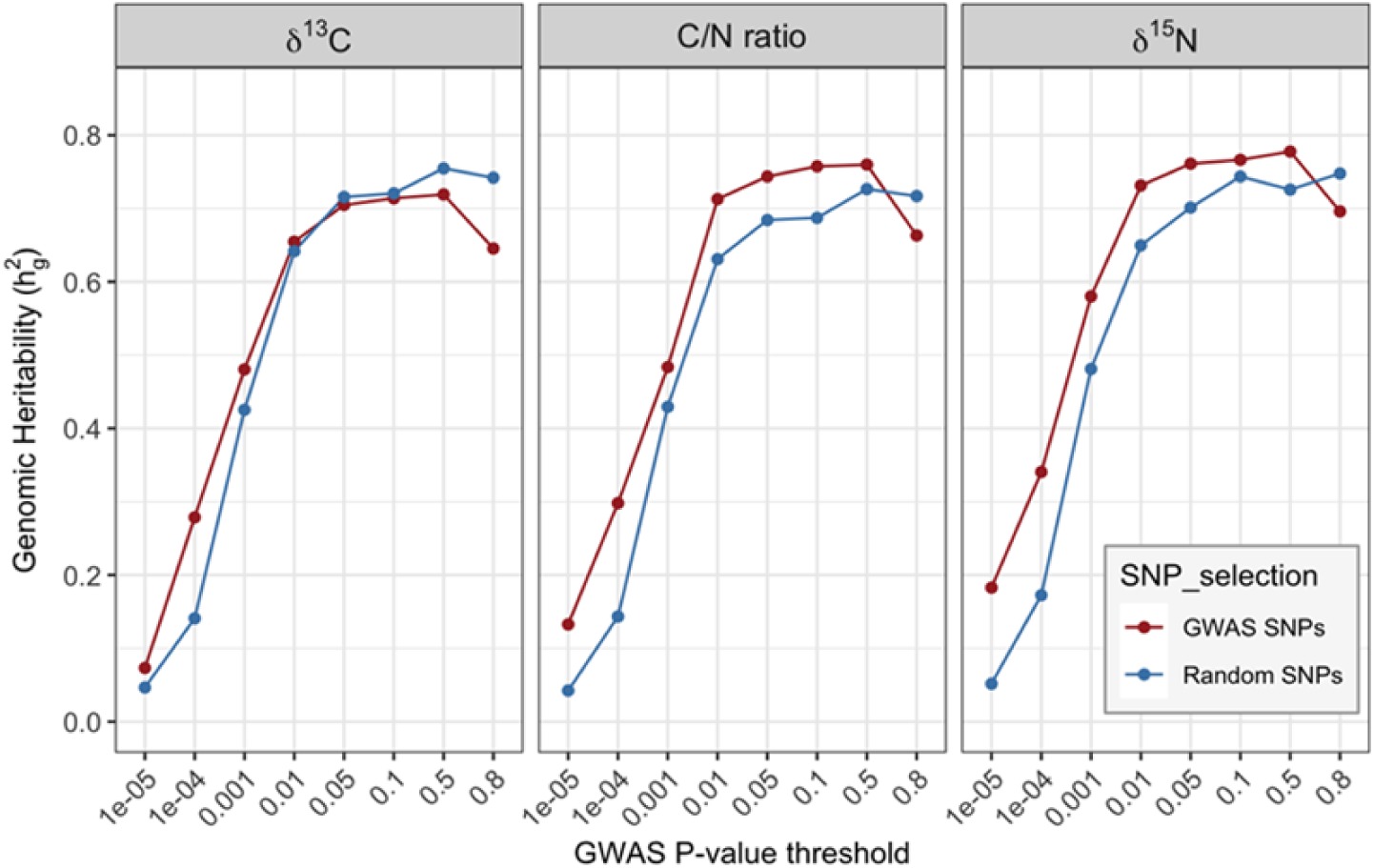
Genomic heritability (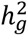) estimates, expressed as the proportion of phenotypic variance explained by SNP markers selected using (i) GWAS P-value significant thresholds and (ii) equally sized random SNP subsets matched to the number of SNPs at each GWAS P-value threshold.

### Genomic Prediction of δ¹³C, δ¹⁵N, and C/N ratio

Within-year genomic prediction ability, using Pearson’s correlation (*r*) between observed phenotypic values and genomic-estimated values, were consistently high across all traits and years for the GBLUP and BRR models (**Figure 3**). δ¹⁵N achieved the highest mean *r* of 0.82 (GBLUP), significantly higher than for δ¹³C (*r* = 0.74) and C/N ratio (*r* = 0.75), which is consistent with their heritability estimates. These prediction results exceed typical values reported for the traits in conifers (*r* = 0.30–0.55; Resende et al., 2017; Isik et al., 2019) and align with the upper range observed for drought-adaptive traits in temperate hardwoods (r = 0.60–0.80; Lstibůrek et al., 2020), underscoring the predictive potential of these foliar traits as selection criteria for climate resilience in *Quercus* species. The substantially lower performance of the LightGBM model (δ¹³C: 0.41; δ¹⁵N: 0.41; C/N ratio: 0.35) could be partly due to model overfitting and/or hyperparameter optimization, a common problem with machine learning methods (Cysneiros-Aragão et al., 2023). The comparable performance and superior computational efficiency of GBLUP relative to BRR prompted its use for subsequent cross-provenance and cross-environment predictions.

**Figure 3.**
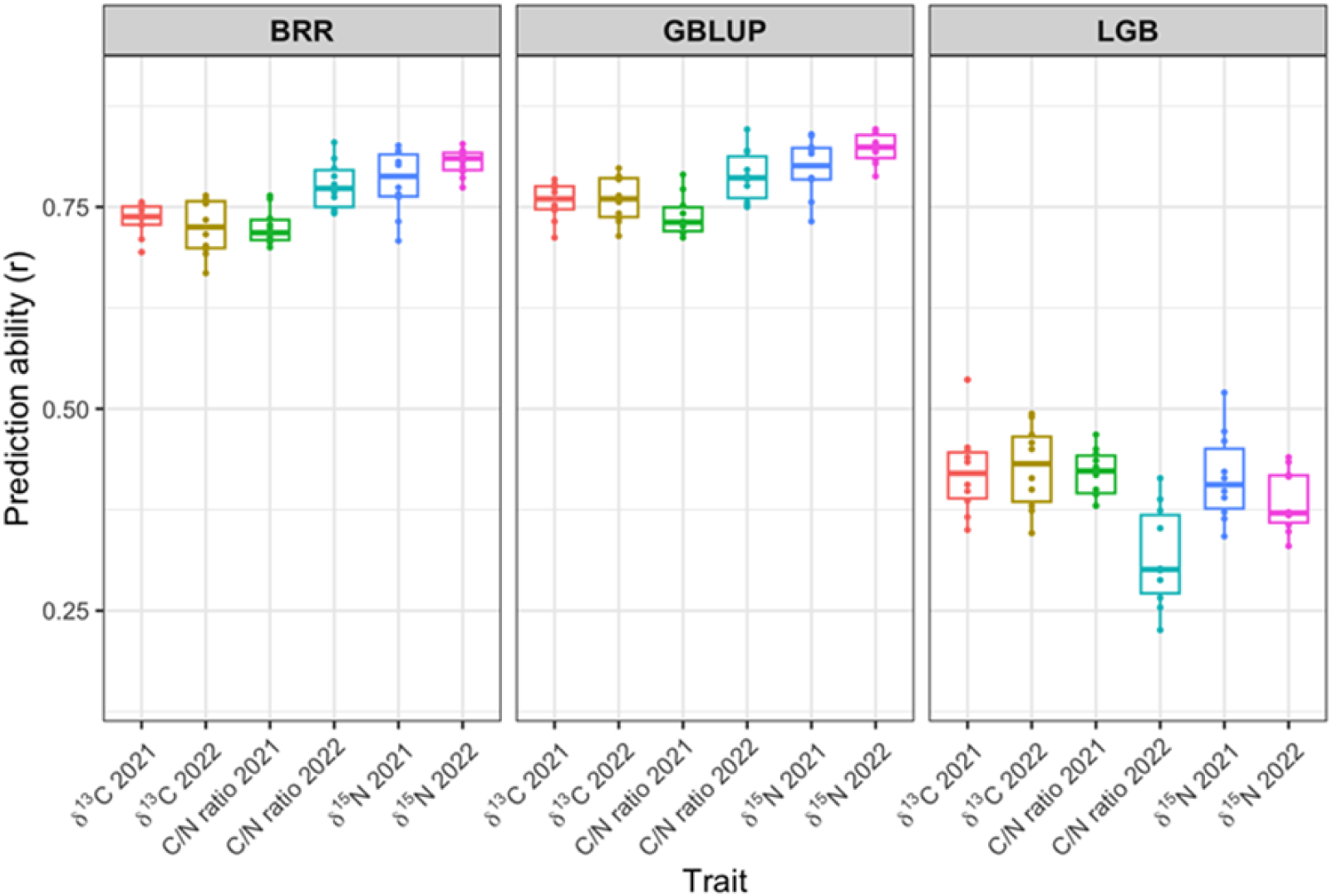
Within-year genomic prediction accuracies for δ¹³C, δ¹⁵N, and C/N ratio and for year 2021 and 2022, obtained with three models (GBLUP, BRR, LGB) using 10-fold cross-validation, with prediction ability quantified as Pearson’s correlation between observed phenotypes and genomic estimated values in the test sets.

### Provenance-Aware Genomic Prediction

Model transferability across provenances is a persistent challenge that often erodes prediction ability in operational forest tree breeding programs (Grattapaglia, 2022). The leave-one-provenance-out predictions yielded robust accuracies (Pearson’s *r*), ranging from 0.54–0.86 for δ¹³C, 0.78–0.92 for δ¹⁵N, and 0.61–0.91 for the C/N ratio, with consistently higher performance in 2022 than in 2021, suggesting improved phenotyping precision or reduced environmental noise in the second sampling year (**Figure 4A**). Notably, the Dymock provenance frequently exhibited reduced predictability across traits and years. The more stringent leave-one-provenance–and-one–year-out cross-validation further demonstrated practical utility, achieving moderate to high prediction abilities for δ¹³C (*r* = 0.41–0.68), δ¹⁵N (*r* = 0.62–0.83), and the C/N ratio (*r* = 0.46–0.91) across novel environment combinations (**Figure 4B**). Although prediction accuracies were lower than those obtained with the leave-one-provenance-out scheme, these results confirm substantial transferability of genomic prediction models for physiological traits in sessile oak, supporting reliable selection of resilient genotypes even when training data originate from distinct provenance–environment contexts.

**Figure 4.**
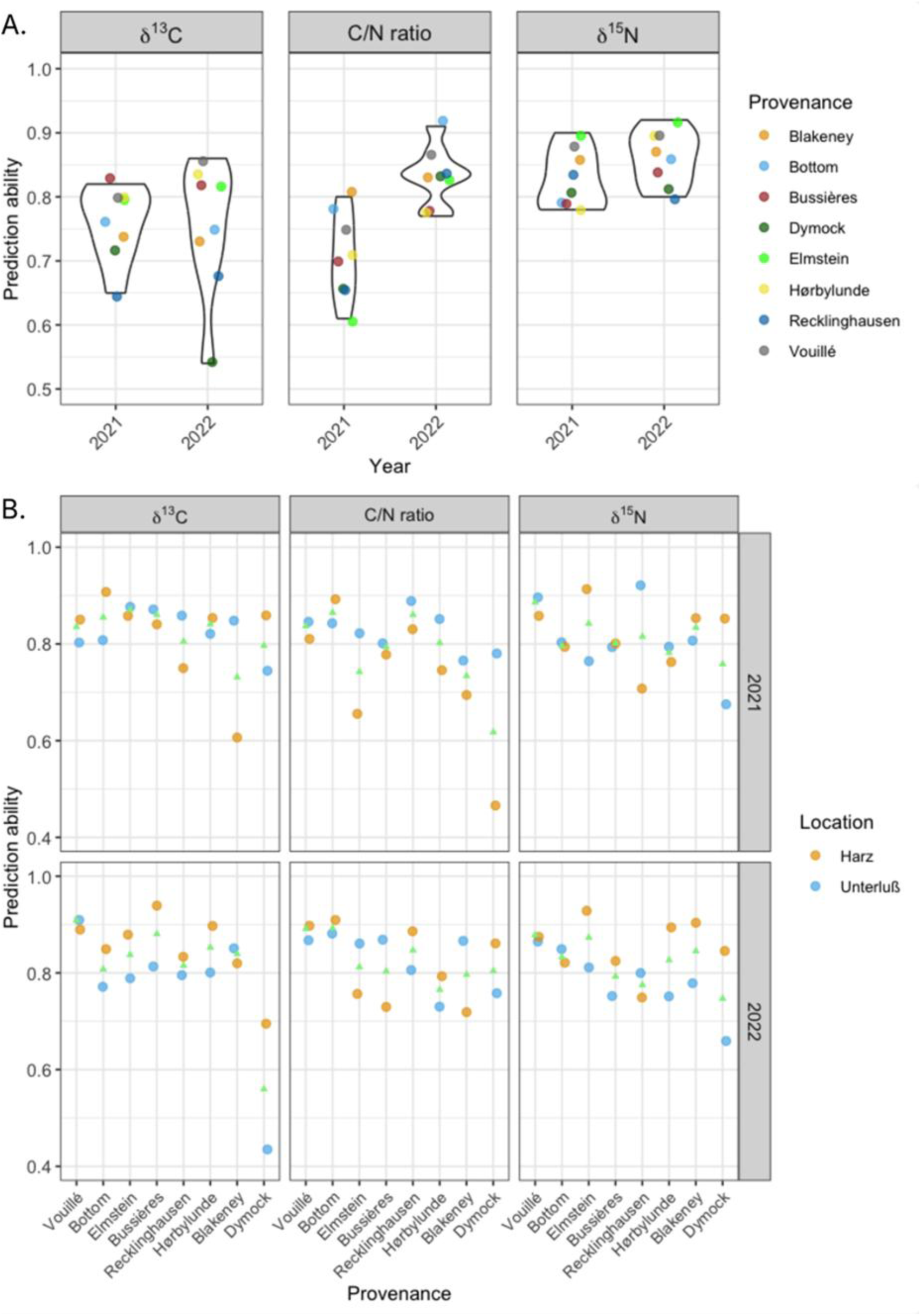
Genomic prediction of each of the eight provenances for each trait using the GBLUP model in (i) the “leave-one-provenance-out” within cross-validation scenario **(A)**, and (ii) the “leave-one-provenance-and-one-year-out” cross-validation scenario **(B)**. Prediction ability measured as Pearson’s correlation between estimated genetic values and observed phenotypic values. Green triangles between points per provenance show the mean prediction ability across the two sites within each year.

### Effect of Genetic Differentiation on Prediction of Provenances

To evaluate how genetic distance influences model transferability, pairwise *F_ST_* values were calculated between each test provenance and the seven training provenances under the leave-one-provenance–and-one-year-out validation scheme. *F_ST_* values ranged from 0 to 0.056 across provenance pairs, indicating moderate population structure among the eight sessile oak origins (Figure 5). Provenance Elmstein (Germany) consistently exhibited the highest *F_ST_* values, yet maintained robust prediction accuracies (mean *r* = 0.87 in 2021; 0.84 in 2022). In contrast, Dymock (United Kingdom), despite showing moderate to high *F_ST_* values (0.016–0.044), displayed substantially reduced prediction ability across traits (mean *r* = 0.72 in 2021; 0.60 in 2022), reflecting its greater geographic separation from the German trial sites (Harz and Unterlüß) and relatively stronger provenance × environment interactions associated with phenotypic plasticity.

**Figure 5.**
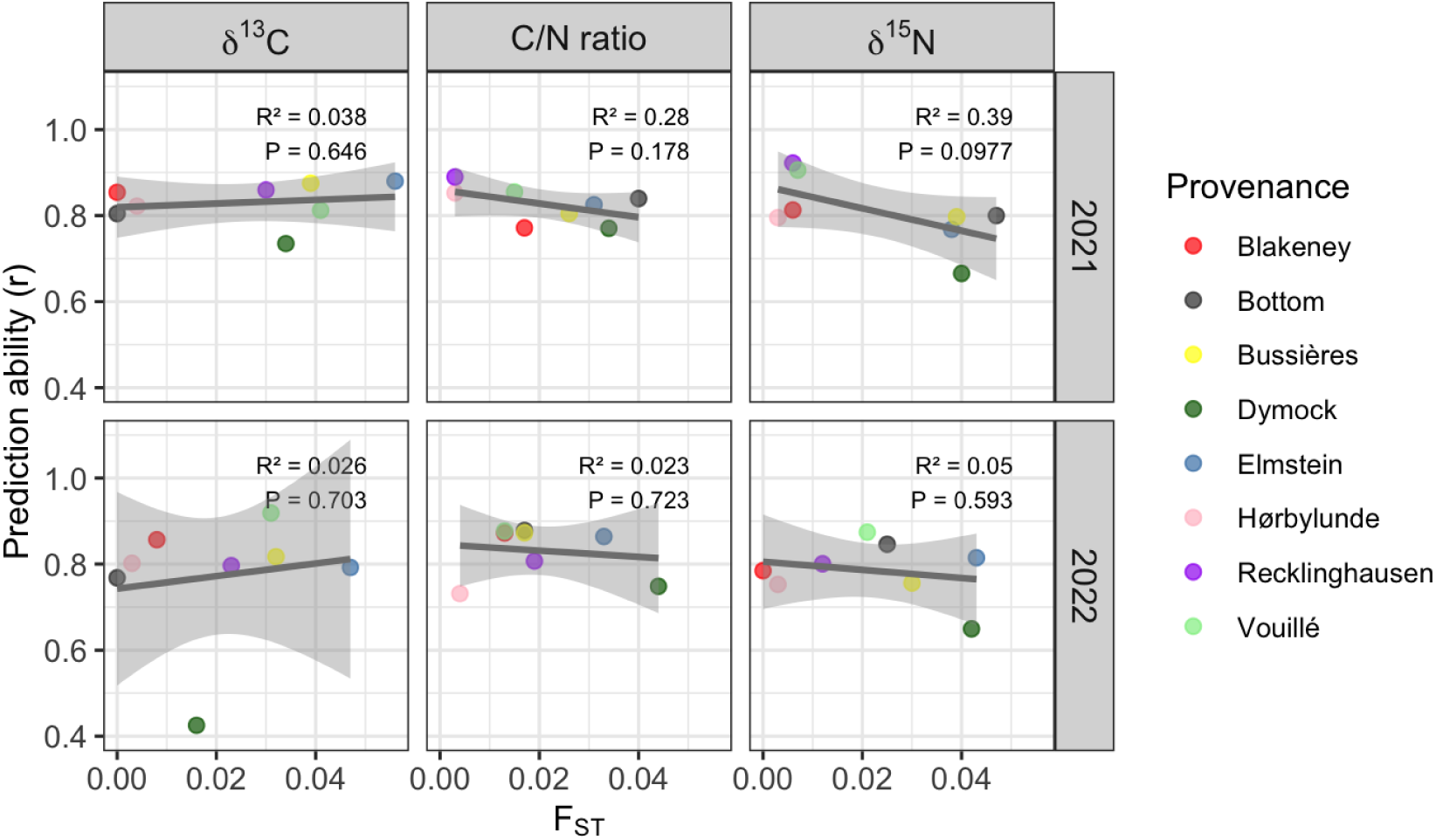
Regression of *F_ST_ between* the tested provenance and the remaining seven training provenances on prediction ability for each trait, illustrates the effect of genetic distance on prediction ability. *F_ST_* were calculated based on GWAS pre-selected SNPs for each trait at a P-value threshold of 0.05.

Prediction accuracy for δ¹⁵N and the C/N ratio declined with increasing *F_ST_*, suggesting genetic distance as one constraint on prediction model transferability. For δ¹³C, there was no relationship with prediction accuracy and *F_ST_*. This pattern for δ¹⁵N and the C/N ratio was more pronounced in 2021, possibly reflecting stronger environmental modulation of genetic signals underlying foliar nitrogen dynamics in year 2021 (Figure 5).

## Discussion

Our results provide one of the first empirical demonstrations of genomic prediction in *Quercus* species for key foliar physiological traits — carbon isotope composition (δ¹³C), nitrogen isotope composition (δ¹⁵N), and the carbon-to-nitrogen ratio (C/N ratio). These traits capture variation in intrinsic water-use efficiency, nitrogen-use efficiency, and soil nitrogen availability (Yang et al., 2015; Nosenko et al., 2025). We obtained high genomic prediction accuracies for δ¹³C (*r* = 0.74), δ¹⁵N (*r* = 0.82), and C/N ratio (*r* = 0.75). These were consistently high across years, confirming the strong phenotypic correlations between the two years. And, further indicate predominant genetic control of these traits, while phenotypic plasticity is reflected in pronounced site effects.

In forest trees, genomic prediction has been extensively investigated for wood and growth traits, with reported accuracies ranging from 0.03 to 0.87 (Duarte et al., 2024; Thistlethwaite et al., 2020; El-Kassaby et al., 2024; Slavov et al., 2025; Akutsu et al., 2023), yet their early evaluation is generally less feasible for operational genomic selection than for physiological traits. Consequently, δ¹³C, δ¹⁵N, and C/N ratio emerge as promising selection targets within a genomic selection framework for climate adaptation, and as early phenotypes for indirect selection on growth-related properties — an approach that is particularly valuable given the increasing frequency of drought and nutrient stress in European forests (Gazol & Camarero, 2022).

Provenance-by-environment interactions (PEI) which were consistently noticeable in δ¹⁵N (Supplementary Figure 6.) marginally reduced prediction accuracy for both δ¹⁵N and C/N ratio. In a meta-analysis encompassing 88 European tree species, Aspalter et al. (2025) reported significant intra-specific PEI effects across several traits, with particularly strong effects in drought-related traits. Li et al. (2015) partly attributed genotype-by-environment variation in growth traits of pine species to site-specific differences in soil nutrient availability, particularly nitrogen and phosphorus. Similar patterns were reported by Lauer et al. (2021), in *Pinus taeda* L., showing that increasing genetic distance between training and test provenances progressively eroded prediction accuracy. In this study, prediction accuracy of δ¹⁵N revealed a negative though non-significant association with *F_ST_* between training and test populations. However, the trend aligns strongly with the pronounced provenance-level differentiation of δ¹⁵N in other studies. δ¹^3^C had no noticeable relationship observed between genetic distance and prediction accuracy. This agrees with Kurjak et al. (2024), showing a weak difference in inter-provenance variation in δ¹³C for *Fagus sylvatica* L. Meanwhile, the number of provenances used in this study is limited; further studies need to explore more provenances with a broader geographic spread to further substantiate these findings.

The high genomic heritability estimates observed in this study (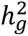 = 0.72–0.78) underscore the strong genetic control of δ¹³C, δ¹⁵N and of C/N ratio in *Quercus petraea*, consistent with reports in the literature. Notably, Alexandre et al. (2020) using observed phenotypic variance, mentioned similar high heritabilities for δ¹³C (up to 0.87) and δ¹⁵N (up to 0.89) in two *Quercus* species. A markedly lower heritability for δ¹³C in *Q. petraea* (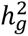 = 0.06) compared to *Q. robur* (h² = 0.87) and a contrasting pattern for C/N ratio (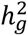 in *Q. petraea* vs. 0.0001 in *Q. robur*) was observed, showing pronounced differences in heritability estimates in closely related oak species (Alexandre et al., 2020). Broad-sense heritability estimates of these traits in other forest trees include 0.48 for δ¹³C, 0.08 for δ¹⁵N, and 0.21 for C/N ratio in *Populus trichocarpa* (McKown et al., 2013); 0.33 for δ¹³C and 0.21 for δ¹⁵N in *Araucaria cunninghamii* (Xu et al., 2003). Collectively, these reports confirm that δ¹³C, δ¹⁵N, and C/N ratio are highly heritable and promising candidates for genomic selection (GS) in forest tree breeding programs. Meanwhile, we further identified a parsimonious genome-wide marker requirement ranging between 30,000 and 50,000 SNPs for maximal extraction of the additive variance present in the studied panel, supporting a high polygenic inheritance of these traits.

GWAS-guided SNP preselection substantially enhanced genomic heritability estimates compared to random sampling, with the largest gains at low-to-moderate marker densities (<50,000 SNPs). Although random SNP subsampling is commonly used for managing dense SNP data in forest tree species (Resende et al., 2017; Grattapaglia et al., 2018), this study demonstrates ∼15% losses in additive variance when using equivalently sized random subsets versus GWAS-prioritized markers. This approach has been used in a few other reports in forest tree species (El-Kassaby et al., 2024; Lobo et al., 2026) and non-tree species (Meuwissen et al., 2024; Izquierdo et al., 2025). It is especially advantageous for outcrossing trees exhibiting high heterozygosity, expansive genomes, and rapid linkage disequilibrium decay (Butler et al., 2022; Plomion et al., 2016).

In our study, prediction performance hinged more on marker density and genetic relatedness between training and test sets than on model choice (GBLUP, BRR, and LightGBM). GBLUP assumes uniform SNP effects (Meuwissen et al., 2001), yet SNPs proximal to causal loci disproportionately drive variance; the BRR model addresses this via variable shrinkage (Habier et al., 2011), yielding simulation gains but modest real-data improvements amid high computational costs (Calus et al., 2016). Here, GWAS preselection positioned the GBLUP model as competitive to other statistical and machine learning methods, thereby streamlining genomic selection pipelines for analytical and computational simplicity with efficiency. The machine learning model (LightGBM) was less precise in predicting phenotypes compared to the GBLUP and BRR models. This may partially result from model overfitting and/or suboptimal hyperparameter tuning, which are common challenges associated with machine learning approaches (Cysneiros-Aragão et al., 2023).

In conclusion, our results highlight the potential of genomic prediction for foliar physiological traits (δ¹³C, δ¹⁵N, and C/N ratio) in forest tree species. High heritability and predictive accuracy position these early-age, easily measured traits — strongly correlated with growth and climate adaptation — as ideal selection targets for breeding climate-resilient forests. Genetic differentiation between training and testing populations may reduce prediction accuracy, particularly for nitrogen-related traits, while GWAS-based SNP pre-selection effectively identified informative markers and revealed polygenic architecture of these traits.

## Acknowledgment

We acknowledge the contribution of Jeremias Götz and Robin Köbel for participating in the sampling exercise in 2021 and 2022, Gesellschaft für Wissenschaftliche Datenverarbeitung GmbH (GWDG), Göttingen, for providing high-performing cluster service for the computational analysis.

## Authors contribution

OG, MM, LVA, WS, HW contributed to the design of the study and methodology. MM and MH^1^ curated the dataset. LVA formally analyzed the data and wrote the original draft of the manuscript. OG, MM, HW reviewed and edited the original manuscript. AH, WS and MH^4^ managed the field trials and reviewed the manuscript. WS, HW, MM, OG acquired funding and were responsible for the project administration. All authors approved the final manuscript.

## Data Availability

Phenotype, genotype, and covariate data used for this study are available at https://figshare.com/s/389883b873489a4e2f90

## Supplementary Figures

**Supplementary Figure 1.**
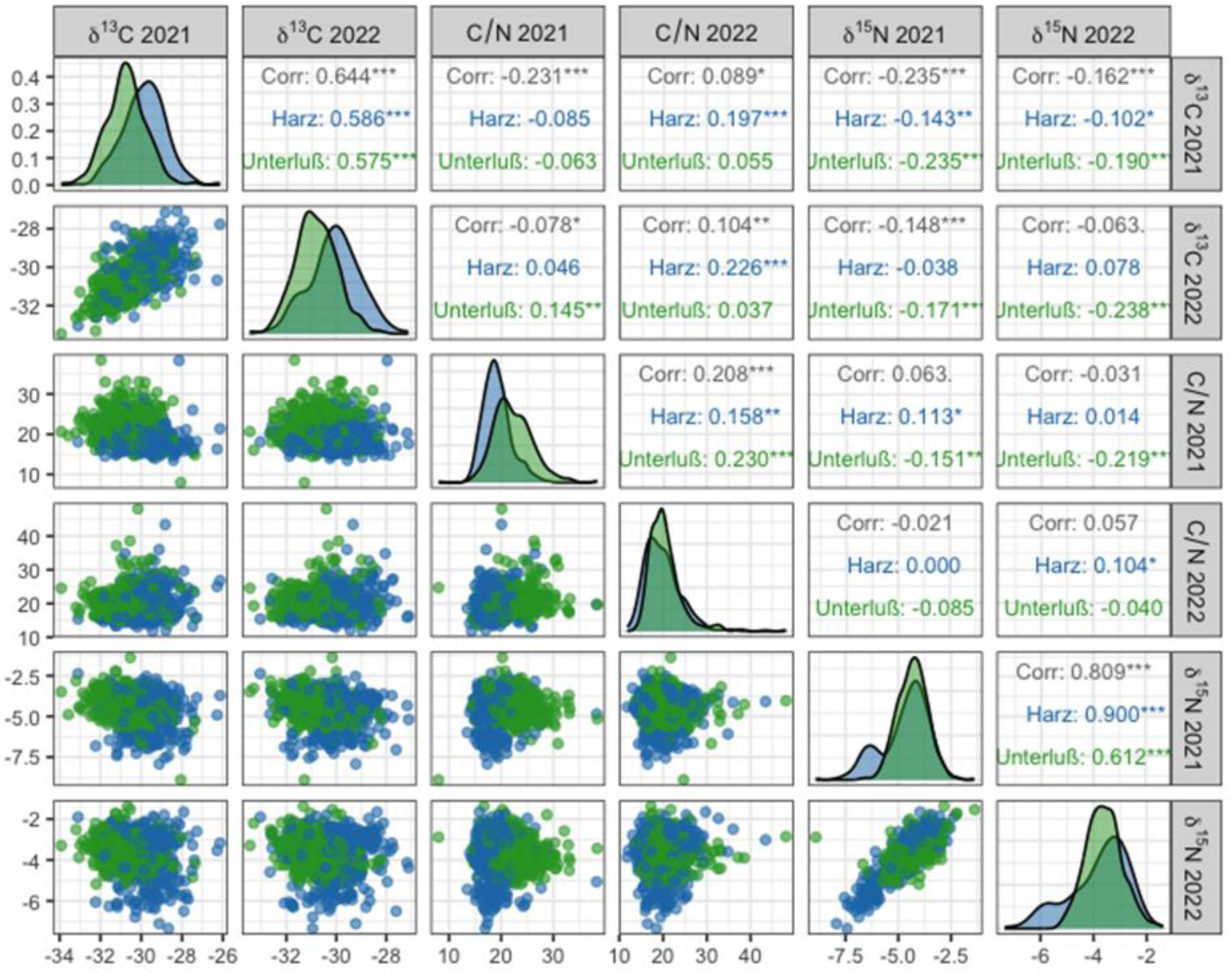
Distributions and pairwise correlations of δ¹³C, δ¹⁵N, and C/N ratio for the Harz and Unterlüß sites in 2021 and 2022, including indications of the statistical significance of each trait correlation. The grey text “corr” shows correlations across sites, while steel blue and green text represent correlations for Harz and Unterlüß sites, respectively.

**Supplementary Figure 2.**
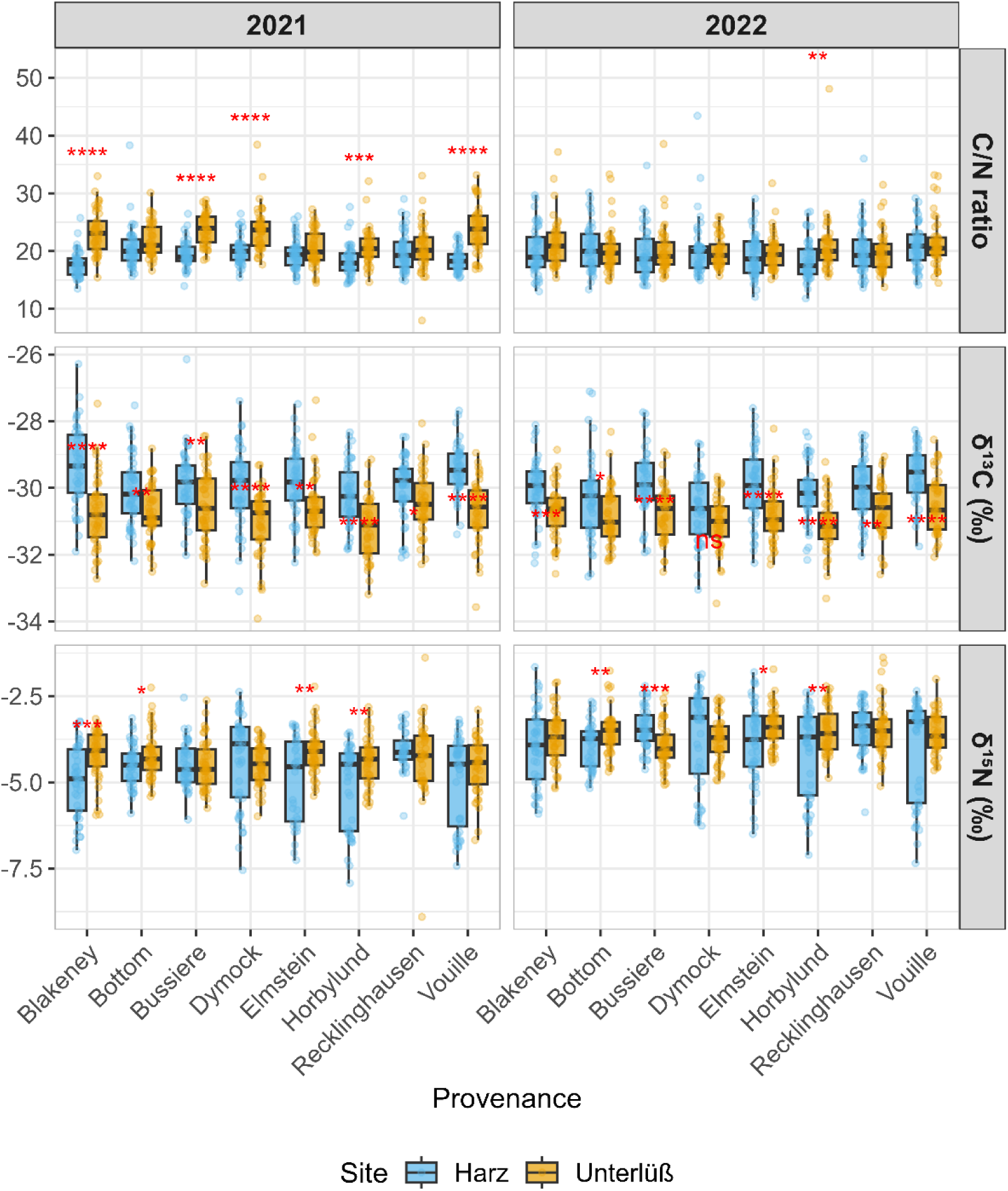
Distributions of δ¹³C, δ¹⁵N, and C/N ratio for provenances within each year (2021 and 2022) and between sites (Harz and Unterlüß), illustrating site effects on trait distribution of each provenance for each trait. The red asterisk shows the provenances with significant site effect at p<0.05.

**Supplementary Figure 3.**
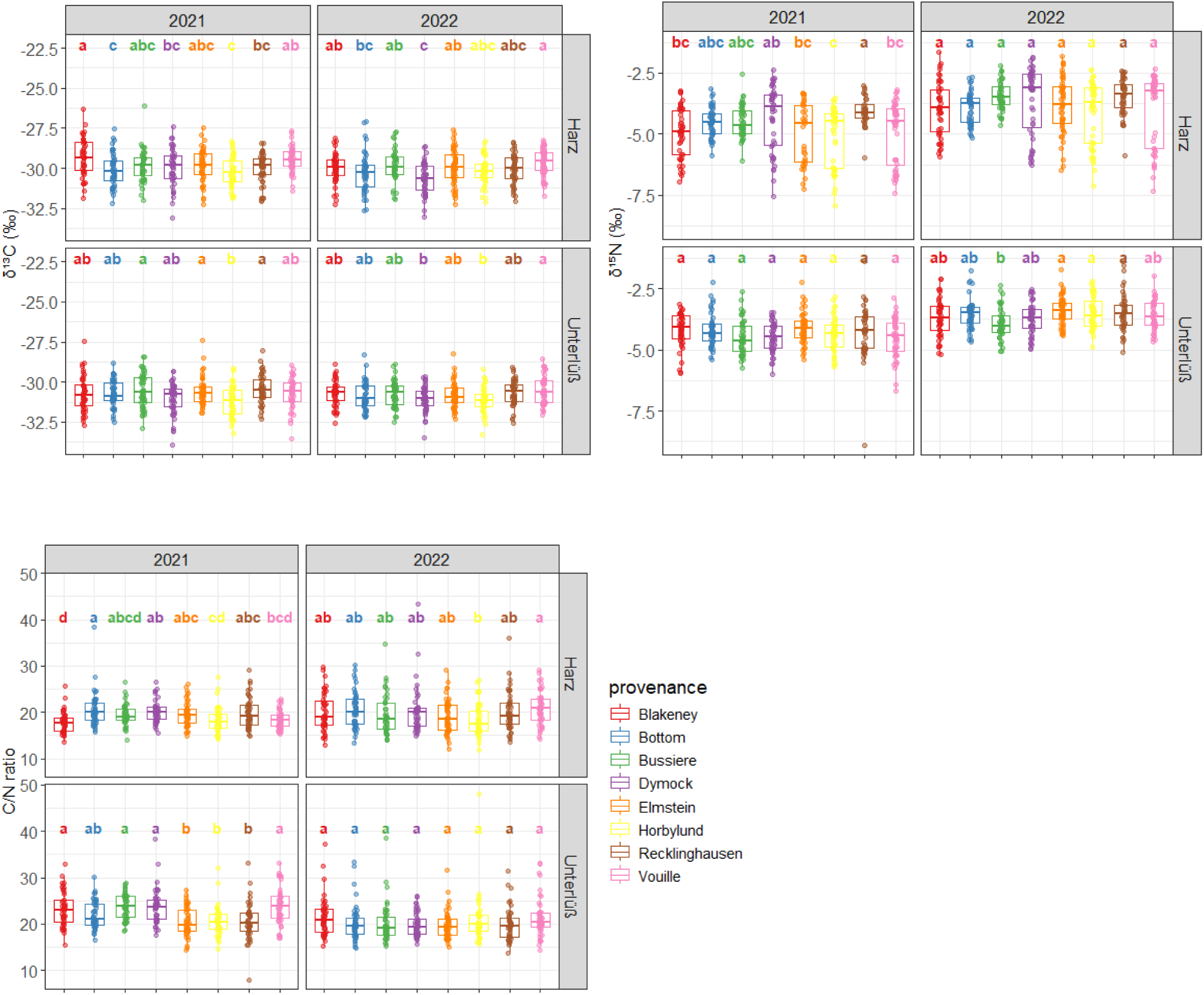
Distributions of δ¹³C, δ¹⁵N, and C/N ratio across provenances within each site (Harz and Unterlüß) and year (2021 and 2022), illustrating differences in mean phenotypic values of each provenance for each trait.

**Supplementary Figure 4.**
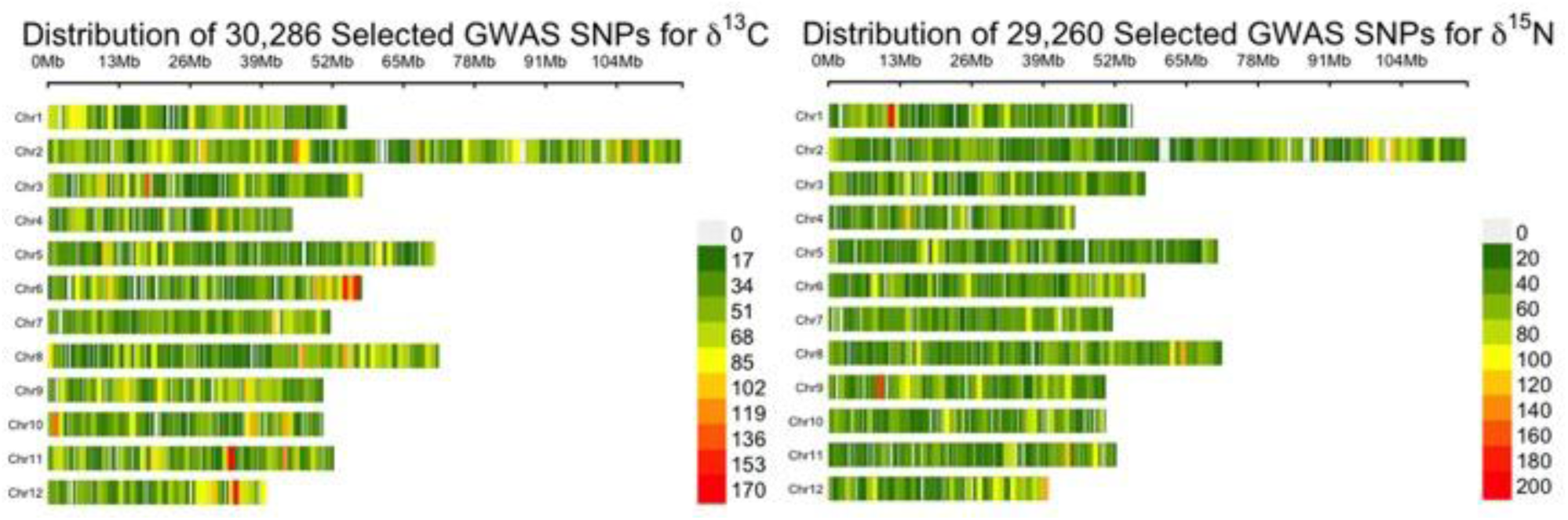
Genome-wide distribution of GWAS-significant SNPs (at P< 0.05) selected for genomic prediction of δ¹³C and δ¹⁵N, respectively.

**Supplementary Figure 5.**
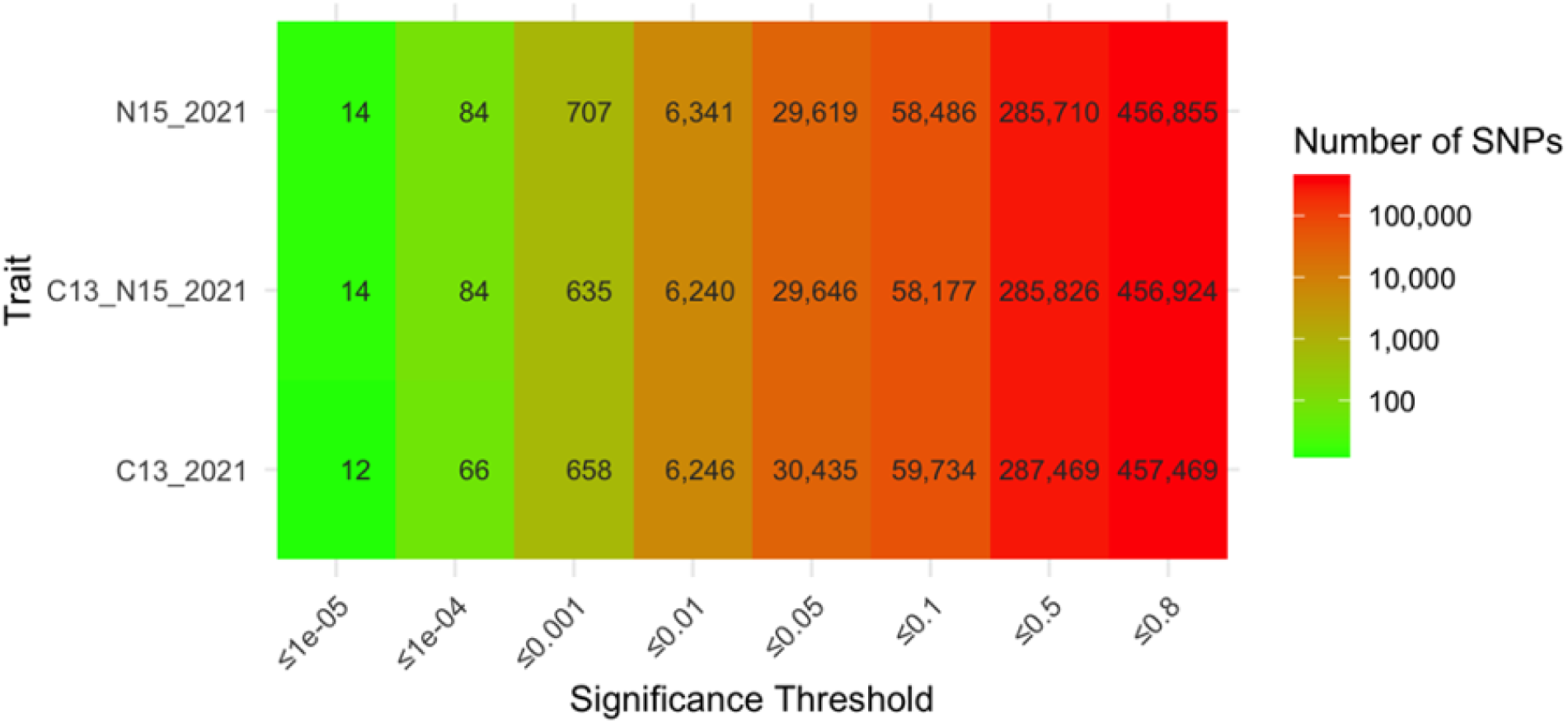
Significance P-value threshold and number of selected GWAS SNPs at each threshold used in genomic heritability (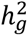) estimation across the three traits.

**Supplementary Figure 6.**
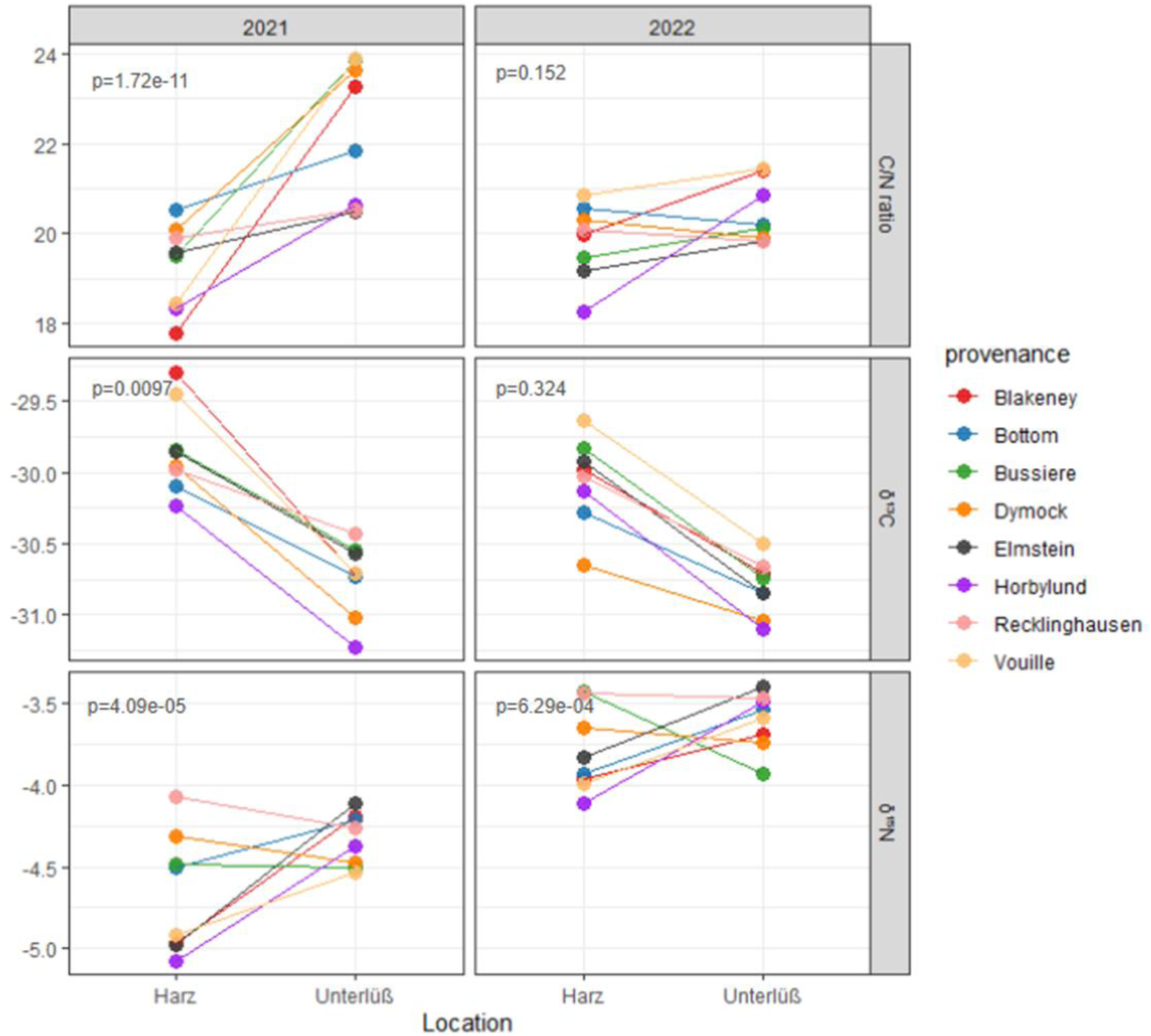
Provenance-by-environment interactions for the eight provenances and between the two sites.

## Supplementary Tables

**Supplementary Table 1.** Climate and geographic data of provenances of origin of the 746 tree samples used in this study. Climate data was retrieved from Wang et al. (2012).

| **Provenance** | **Country** | **Lat (°N)** | **Long (°E)** | **Ele (masl)** | **MAT** | **MWMT** | **MCMT** | **TD** | **MAP** | **MSP** |
| --- | --- | --- | --- | --- | --- | --- | --- | --- | --- | --- |
| Hørbylund | Denmark | 56° 08' N | 9° 25' E | 77 | 7.2 | 15.5 | -1.4 | 16.9 | 765 | 331 |
| Elmstein-Nord | Germany | 49° 30' N | 7° 34' E | 450 | 7.7 | 16.2 | -1.3 | 17.5 | 729 | 339 |
| Recklinghausen | Germany | 51° 46' N | 7° 11' E | 75 | 9.4 | 17.2 | 1.5 | 15.7 | 815 | 365 |
| Bussieres | France | 47° 45' N | 5° 32' E | 360 | 9.4 | 17.9 | 0.6 | 17.3 | 892 | 364 |
| Voûille St. Hilaire | France | 46° 38' N | 0° 10' W | 143 | 11.2 | 18.8 | 3.9 | 14.9 | 786 | 289 |
| Dean Blakeney | Great Britain | 51° 47' N | 2° 30' W | 76 | 9.6 | 16.1 | 3.9 | 12.2 | 806 | 318 |
| Dean Sutton Bottom | Great Britain | 51° 48' N | 2° 29' W | 120 | 9.4 | 16.0 | 3.6 | 12.4 | 800 | 317 |
| Dymock | Great Britain | 51° 57' N | 2° 27' W | 70 | 9.6 | 16.1 | 3.9 | 12.2 | 717 | 296 |

\*Lat - latitude, \*Long - longitude, \*Ele - elevation, \*MAT - mean annual temperature, \*MWMT - mean warmest month temperature, \*MCMT – mean coldest month temperature, \*TD - temperature difference (mean annual), \*MAP - mean annual precipitation, \*MSP - mean summer precipitation

**Supplementary Table 2.** Statistical results for site effect on each provenance for δ¹³C, δ¹⁵N, and C/N ratio.

| Provenance | Trait | Year | No. of trees<br>in site Harz | No of tree in<br>site<br>Unterlüß | P-value | Adjusted P-value (fdr) | Statistical<br>Significance |
| --- | --- | --- | --- | --- | --- | --- | --- |
| **Blakeney** | C/N ratio | 2021 | 46 | 46 | 1.02e-12 | 3.31e-08 | **** |
| **Blakeney** | C/N ratio | 2022 | 46 | 45 | 0.101 | 0.1426 | ns |
| **Blakeney** | $\delta^{13}\text{C}$ | 2021 | 46 | 46 | 3.58e-08 | 3.44e-03 | **** |
| **Blakeney** | $\delta^{13}\text{C}$ | 2022 | 46 | 45 | 7.89e-05 | 0.0003 | *** |
| **Blakeney** | $\delta^{15}\text{N}$ | 2021 | 46 | 46 | 6.00e-05 | 0.0002 | *** |
| **Blakeney** | $\delta^{15}\text{N}$ | 2022 | 46 | 45 | 0.164 | 0.2187 | ns |
| **Bottom** | C/N ratio | 2021 | 46 | 48 | 0.0548 | 0.0849 | ns |
| **Bottom** | C/N ratio | 2022 | 46 | 48 | 0.645 | 0.7036 | ns |
| **Bottom** | $\delta^{13}\text{C}$ | 2021 | 46 | 48 | 0.00127 | 0.0029 | ** |
| **Bottom** | $\delta^{13}\text{C}$ | 2022 | 45 | 48 | 0.0128 | 0.0236 | * |
| **Bottom** | $\delta^{15}\text{N}$ | 2021 | 46 | 48 | 0.0243 | 0.0402 | * |
| **Bottom** | $\delta^{15}\text{N}$ | 2022 | 45 | 48 | 0.00343 | 0.0072 | ** |
| **Bussiere** | C/N ratio | 2021 | 44 | 47 | 1.38e-12 | 3.31e-08 | **** |
| **Bussiere** | C/N ratio | 2022 | 44 | 46 | 0.461 | 0.5406 | ns |
| **Bussiere** | $\delta^{13}\text{C}$ | 2021 | 44 | 47 | 0.00182 | 0.0040 | ** |
| **Bussiere** | $\delta^{13}\text{C}$ | 2022 | 44 | 46 | 7.77e-06 | 3.39e-09 | **** |
| **Bussiere** | $\delta^{15}\text{N}$ | 2021 | 44 | 47 | 0.886 | 0.887 | ns |
| **Bussiere** | $\delta^{15}\text{N}$ | 2022 | 44 | 46 | 0.000218 | 0.000669 | *** |
| **Dymock** | C/N ratio | 2021 | 46 | 48 | 3.01e-07 | 2.16e-06 | **** |
| **Dymock** | C/N ratio | 2022 | 46 | 46 | 0.625 | 0.6977 | ns |
| **Dymock** | $\delta^{13}\text{C}$ | 2021 | 46 | 48 | 7.01e-06 | 3.39e-09 | **** |
| **Dymock** | $\delta^{13}\text{C}$ | 2022 | 46 | 46 | 0.0358 | 0.05728 | ns |
| **Dymock** | $\delta^{15}\text{N}$ | 2021 | 46 | 48 | 0.473 | 0.5406 | ns |
| **Dymock** | $\delta^{15}\text{N}$ | 2022 | 46 | 46 | 0.705 | 0.752 | ns |
| **Elmstein** | C/N ratio | 2021 | 48 | 48 | 0.115 | 0.1577 | ns |
| **Elmstein** | C/N ratio | 2022 | 48 | 48 | 0.336 | 0.4244 | ns |
| **Elmstein** | $\delta^{13}\text{C}$ | 2021 | 48 | 48 | 0.000362 | 0.0010 | ** |
| **Elmstein** | $\delta^{13}\text{C}$ | 2022 | 48 | 48 | 1.03e-05 | 4.12e-05 | **** |
| **Elmstein** | $\delta^{15}\text{N}$ | 2021 | 33 | 48 | 0.000808 | 0.0020 | ** |
| **Elmstein** | $\delta^{15}\text{N}$ | 2022 | 48 | 48 | 0.0208 | 0.0357 | * |
| **Horbylund** | C/N ratio | 2021 | 47 | 45 | 0.000223 | 0.000669 | *** |
| **Horbylund** | C/N ratio | 2022 | 46 | 45 | 0.00475 | 0.0094848 | ** |
| **Horbylund** | $\delta^{13}\text{C}$ | 2021 | 47 | 45 | 1.36e-06 | 8.16e-06 | **** |
| **Horbylund** | $\delta^{13}\text{C}$ | 2022 | 46 | 45 | 3.15e-07 | 2.16e-06 | **** |
| **Horbylund** | $\delta^{15}\text{N}$ | 2021 | 47 | 45 | 0.00119 | 0.002856 | ** |
| **Horbylund** | $\delta^{15}\text{N}$ | 2022 | 47 | 45 | 0.00494 | 0.0094848 | ** |
| **Recklinghausen** | C/N ratio | 2021 | 48 | 45 | 0.41 | 0.5046 | ns |
| **Recklinghausen** | C/N ratio | 2022 | 47 | 45 | 0.757 | 0.7899 | ns |
| **Recklinghausen** | $\delta^{13}\text{C}$ | 2021 | 48 | 45 | 0.0188 | 0.0334 | * |
| **Recklinghausen** | $\delta^{13}\text{C}$ | 2022 | 47 | 45 | 0.000646 | 0.0017 | ** |
| **Recklinghausen** | $\delta^{15}\text{N}$ | 2021 | 33 | 45 | 0.314 | 0.4074 | ns |
| **Recklinghausen** | $\delta^{15}\text{N}$ | 2022 | 47 | 45 | 0.887 | 0.887 | ns |
| **Vouille** | C/N ratio | 2021 | 48 | 45 | 1.94e-11 | 3.10e-07 | **** |
| **Vouille** | C/N ratio | 2022 | 48 | 44 | 0.472 | 0.5406 | ns |
| **Vouille** | $\delta^{13}\text{C}$ | 2021 | 48 | 45 | 4.46e-10 | 5.35e-06 | **** |
| **Vouille** | $\delta^{13}\text{C}$ | 2022 | 48 | 44 | 7.36e-06 | 3.39e-09 | **** |
| **Vouille** | $\delta^{15}\text{N}$ | 2021 | 48 | 45 | 0.0849 | 0.1260 | ns |
| **Vouille** | $\delta^{15}\text{N}$ | 2022 | 48 | 44 | 0.0866 | 0.1260 | ns |

Significance codes: '\*\*\*\*' p<0.0001; '\*\*\*' p<0.001; '\*\*' p<0.01; '\*' p<0.05; ns - not significant.

**Supplementary Table 3.** Analysis of variance showing provenance, site, and provenance x site effects on δ¹³C, δ¹⁵N, and C/N ratio.

| $\delta^{13}\text{C}$ 2021 | Df | SS | MS | F-value | P-value | Signif | % SS |
| --- | --- | --- | --- | --- | --- | --- | --- |
| **Site** | 1 | 152.3 | 152.26 | 159.251 | 2.00e-16 | *** | 16.85 |
| **provenance** | 7 | 36.4 | 5.21 | 5.445 | 4.15e-06 | *** | 4.03 |
| **Site x provenance** | 7 | 17.9 | 2.55 | 2.672 | 0.00979 | ** | 1.98 |
| **Residuals** | 729 | 697.0 | 0.96 |  |  |  | 77.14 |
| $\delta^{13}\text{C}$ 2022 | | | | | | | |
| **Site** | 1 | 105.0 | 105.05 | 127.428 | 2.00e-16 | *** | 14.16 |
| **provenance** | 7 | 36.0 | 5.14 | 6.239 | 4.08e-07 | *** | 4.85 |
| **Site x provenance** | 7 | 6.7 | 0.96 | 1.160 | 0.324 |  | 0.90 |
| **Residuals** | 721 | 594.4 | 0.82 |  |  |  | 80.09 |
| $\delta^{15}\text{N}$ 2021 | | | | | | | |
| **Site** | 1 | 21.5 | 21.474 | 26.131 | 4.12e-07 | *** | 3.34 |
| **provenance** | 7 | 19.4 | 2.770 | 3.370 | 0.00152 | ** | 3.02 |
| **Site x provenance** | 7 | 26.8 | 3.828 | 4.658 | 4.09e-05 | *** | 4.17 |
| **Residuals** | 699 | 574.4 | 0.822 |  |  |  | 89.46 |
| $\delta^{15}\text{N}$ 2022 | | | | | | | |
| **Site** | 1 | 7.0 | 6.993 | 8.441 | 0.003780 | ** | 1.10 |
| **provenance** | 7 | 9.8 | 1.396 | 1.685 | 0.109517 |  | 1.54 |
| **Site x provenance** | 7 | 21.4 | 3.056 | 3.689 | 0.000629 | *** | 3.36 |
| **Residuals** | 722 | 598.1 | 0.828 |  |  |  | 94.00 |
| C/N ratio 2021 |  |  |  |  |  |  |  |
| **Site** | 1 | 1691 | 1690.6 | 178.533 | 2.00e-16 | *** | 17.45 |
| **provenance** | 7 | 454 | 64.8 | 6.845 | 6.79e-08 | *** | 4.68 |
| **Site x provenance** | 7 | 639 | 91.2 | 9.633 | 1.72e-11 | *** | 6.59 |
| **Residuals** | 729 | 6903 | 9.5 |  |  |  | 71.28 |
| C/N ratio 2022 |  |  |  |  |  |  |  |
| **Site** | 1 | 66 | 65.80 | 4.237 | 0.0399 | * | 0.56 |
| **provenance** | 7 | 213 | 30.39 | 1.957 | 0.0584 | . | 1.82 |
| **Site x provenance** | 7 | 167 | 23.84 | 1.535 | 0.1521 |  | 1.43 |
| **Residuals** | 722 | 11212 | 15.53 |  |  |  | 96.18 |

Significance codes: '\*\*\*' $p < 0.001$ ; '\*\*' $p < 0.01$ ; '\*' $p < 0.05$ ; '.' $P < 0.1$ ; ' ' $p < 1$ ; SS – sum of squares; MS – mean squares; %SS percentage of sum of squares.

**Supplementary Table 4a.** Environmental data for site Harz during the sampling period for year 2021 and 2022.

| Provenance | 2021<br>sampling<br>date | Temp.<br>2021<br>(°C) | Rel.<br>Hum.<br>2021 (%) | 2022<br>sampling<br>date | Temp.<br>2022<br>(°C) | Rel. Hum.<br>2022 (%) |
| --- | --- | --- | --- | --- | --- | --- |
| Voillé, Frankreich | 16/07 | 20.9 | 82.8 | 01/06 | 14.8 | 66.3 |
| Voillé, Frankreich | 16/07 | 22.5 | 77.0 | 02/06 | 14.8 | 67.0 |
| Horbylunde, Denmark | 25/08 | 22.5 | 47.0 | 02/06 | 15.0 | 63.2 |
| Horbylunde, Denmark | 22/07 | 20.8 | 70.2 | 02/06 | 23.7 | 37.7 |
| Horbylunde, Denmark | 22/07 | 19.7 | 67.3 | 01/06 | 15.5 | 57.3 |
| Dean Blackney, Great Britain | 15/07 | 24.9 | 65.3 | 01/06 | 14.9 | 64.0 |
| Dean Blackney, Great Britain | 16/07 | 24.0 | 73.7 | 01/06 | 15.1 | 65.8 |
| Dean Blackney, Great Britain | 15/07 | 24.1 | 71.2 | 02/06 | 19.6 | 45.4 |
| Elmstein-Nord, Germany | 27/07 | 20.6 | 77.6 | 01/06 | NA | NA |
| Elmstein-Nord, Germany | 28/07 | 20.7 | 73.1 | 01/06 | 14.1 | 81.9 |
| Elmstein-Nord, Germany | 13/08 | N/A | N/A | 02/06 | 15.9 | 64.0 |
| Recklinghausen, Germany | 27/07 | 25.0 | 54.9 | 01/06 | 16.6 | 56.0 |
| Recklinghausen, Germany | 13/08 | N/A | N/A | 02/06 | 17.6 | 56.8 |
| Recklinghausen, Germany | N/A | N/A | N/A | 02/06 | 17.6 | 56.8 |
| Recklinghausen, Germany | 29/07 | 18.0 | 71.9 | 15/06 | NA | NA |
| Bussieres, France | 27/07 | 22.4 | 68.5 | 01/06 | 17.4 | 52.0 |
| Bussieres, France | 28/07 | 22.0 | 75.5 | 01/06 | 14.5 | 75.9 |
| Bussieres, France | 29/07 | 18.9 | 67.9 | 02/06 | 20.3 | 43.3 |
| Bussieres, France | N/A | N/A | N/A | 02/06 | 20.3 | 43.3 |
| Dean Sutton Bottow, Great Britain | 28/07 | 20.1 | 75.7 | 01/06 | 17.8 | 53.1 |
| Dean Sutton Bottow, Great Britain | 16/08 | 21.7 | 56.2 | 15/06 | 23.9 | 38.5 |
| Dean Sutton Bottow, Great Britain | N/A | N/A | N/A | 15/06 | 25.3 | 36.0 |
| Dymock, Great Britain | 29/07 | 20.7 | 67.0 | 15/06 | 25.6 | 37.2 |
| Dymock, Great Britain | 13/08 | N/A | N/A | 15/06 | NA | NA |
| Dymock, Great Britain | N/A | N/A | N/A | 15/06 | NA | NA |
| Dymock, Great Britain | 28/07 | 21.5 | 68.7 | 15/06 | 20.7 | 45.6 |

**Supplementary Table 4b.** Environmental data for site Unterlüß during the sampling period for year 2021 and 2022.

| Provenance | 2021<br>sampling<br>date | Temp.<br>2021<br>(°C) | Rel.<br>Hum.<br>2021 (%) | 2022<br>sampling<br>date | Temp.<br>2022<br>(°C) | Rel. Hum.<br>2022 (%) |
| --- | --- | --- | --- | --- | --- | --- |
| Bussieres, France | 25/08 | 9.0 | 57.2 | 20/06 | 22.3 | 49.4 |
| Bussieres, France | 30/08 | 17.2 | 79.7 | 21/06 | 17.8 | 59.5 |
| Bussieres, France | 03/09 | 25.0 | 50.0 | 21/06 | 23.2 | 39.4 |
| Dean Blackney, Great Britain | 25/08 | 16.8 | 70.2 | 20/06 | NA | NA |
| Dean Blackney, Great Britain | 31/08 | 13.4 | 71.2 | 21/06 | 19.8 | 48.4 |
| Dean Blackney, Great Britain | 03/09 | 18.8 | 74.2 | 21/06 | NA | NA |
| Voillé, Frankreich | 25/08 | 17.7 | 74.2 | 20/06 | 17.8 | 60.3 |
| Voillé, Frankreich | 31/08 | 17.0 | 82.5 | 21/06 | NA | NA |
| Voillé, Frankreich | 03/09 | 25.1 | 49.6 | 21/06 | 23.9 | 38.3 |
| Dean Sutton Bottow, Great Britain | 25/08 | 19.5 | 64.9 | 20/06 | 19.4 | 57.2 |
| Dean Sutton Bottow, Great Britain | 30/08 | 17.0 | 84.4 | 21/06 | 17.8 | 60.8 |
| Dean Sutton Bottow, Great Britain | 04/09 | 15.7 | 68.2 | 21/06 | NA | NA |
| Dymock, Great Britain | 25/08 | 10.1 | 70.9 | 20/06 | 16.4 | 60.2 |
| Dymock, Great Britain | 31/08 | 19.4 | 73.5 | 21/06 | 21.5 | 46.4 |
| Dymock, Great Britain | 03/09 | 22.5 | 62.2 | 21/06 | NA | NA |
| Recklinghausen, Germany | 25/08 | 18.7 | 69.2 | 20/06 | 18.0 | 59.0 |
| Recklinghausen, Germany | 31/08 | 16.5 | 82.8 | 21/06 | 19.5 | 52.0 |
| Recklinghausen, Germany | 03/09 | 21.2 | 61.1 | 21/06 | NA | NA |
| Elmstein-Nord, Germany | 30/08 | 17.7 | 81.2 | 20/06 | 17.8 | 55.2 |
| Elmstein-Nord, Germany | 30/08 | 18.8 | 77.0 | 21/06 | 20.2 | 56.2 |
| Elmstein-Nord, Germany | 03/09 | 18.8 | 74.2 | 21/06 | NA | NA |
| Horbylunde, Denmark | 30/08 | 19.0 | 74.0 | 20/06 | NA | NA |
| Horbylunde, Denmark | 31/08 | 18.8 | 76.1 | 21/06 | NA | NA |
| Horbylunde, Denmark | 03/09 | 21.4 | 63.2 | 21/06 | 22.2 | 42.7 |

